# Phylogenetic alpha and beta diversity of African ferns

**DOI:** 10.64898/2026.11.12.766204

**Authors:** Israel T. Borokini, Immaculate K. Constantine, Mwihaki J. Karichu, Susan Fawcett, Joel H. Nitta

**Affiliations:** Department of Ecology, Montana State University Bozeman, 310 Lewis Hall, Bozeman MT 59717; National Herbarium of Tanzania, Tanzania Plant Health and Pesticides Authority (TPHPA), P. O. Box 3024, Arusha, Tanzania; New York Botanical Garden, 2900 Southern Blvd, Bronx, NY 10458; Graduate Center, City University of New York, 365 5th Ave, New York, NY 10016, USA; University and Jepson Herbaria, University of California Berkeley, Berkeley CA 94720; Graduate School of Global and Transdisciplinary Studies, Chiba University, Japan

**Keywords:** phylodiversity, ferns, Africa, endemism

## Abstract

Ferns are widely distributed across nearly all major biomes and are indicators of ecosystem health and functioning. Therefore, their diversity and distributions can be used to elucidate the ecological and evolutionary mechanisms associated with net diversification and deep biogeographical events. Though Africa has relatively low fern species richness, the spatial patterns of fern diversity and distribution is non-random and may be indicative of the evolutionary history and paleo-ecological conditions that shape current floristic diversity on the continent. This study assessed spatial patterns of taxonomic and phylogenetic diversity of ferns in Africa. Using species occurrence records from biodiversity databases (Global Biodiversity Information Facility, the African Plants Portal, RAINBIO, and recently digitized specimens across 31 U.S. herbaria) and the Fern Tree of Life, we calculated and mapped several biodiversity measures including species richness, phylogenetic diversity (PD) and endemism (PE), relative phylogenetic diversity (RPD) and endemism (RPE), Sorensen-based phylogenetic and taxonomic beta diversity, and grade-of-membership models using phylogenetic and taxonomic distances. We also decomposed PE and RPE significance into different endemism centers using the categorical analysis of neo- and paleo-endemism (CANAPE) and investigated lineages driving patterns of endemism with the Range-weighted branch length difference metric (RWiBaLD). Finally, we investigate the association of contemporary climatic and soil factors and topography on spatial phylodiversity patterns of African ferns.

## Introduction

Ferns diverged from seed plants sometime during the Devonian–Carboniferous period, approximately 360–450 million years ago (Pryer *et al*. 2004; Rothwell and Stockey 2008; Nitta *et al*. 2022). Globally, there are approximately 12,000–13,000 species (Pteridophyte Phylogeny Group I 2016), with the highest diversity concentrated in tropical and subtropical regions, particularly in Southeast Asia, the Neotropics (especially the Andes and Amazon Basin), and parts of Oceania (Suissa *et al*. 2021). Compared to these global hotspots, Africa supports a relatively low but ecologically significant diversity of ferns, estimated at 1,200 species across the continent (Aldasoro *et al*. 2004; Constantine *et al*. 2026). Despite this lower species richness, African ferns occupy a wide range of ecological niches, highlighting the continent’s importance for understanding patterns of plant diversity and evolution (Aldasoro *et al*. 2004; Kreft and Jetz 2007; Suissa *et al*. 2021).

Fern diversity in Africa is unevenly distributed, reflecting strong environmental and historical gradients across the continent (Aldasoro *et al*. 2004). The highest species richness occurs in the Guineo-Congolian tropical rainforest, the Eastern Arc mountains and adjacent East African highlands, and Madagascar, where high moisture availability and complex topography create favorable microhabitats (Aldasoro *et al*. 2004; Figs. S1, S2). Consequently, Madagascar is recognized as one of eight global fern diversity hotspots, with secondary areas of high diversity centered in the lower Guinean forest ecosystem and the Mediterranean Cape region (Suissa *et al*. 2021). In contrast, arid and semi-arid regions, including the Sahara and Sahel, exhibit very low fern diversity due to limited water availability and the absence of stable refugia (Brito *et al*. 2014). Beyond the role of climate, elevation, and historical refugia in shaping the spatial distribution of fern diversity across Africa, biological factors such as their reliance on water for reproduction, shade tolerance, and spore-based dispersal, may favor their colonization and persistence in humid and forested environments, while limiting their occurrence in dry habitats (Smith 1993; Aldasoro *et al*. 2004).

Following these spatial patterns of diversity, phylogenetic patterns of African ferns are shaped by a combination of historical diversification, climatic stability, and extinction dynamics (Pryer *et al*. 2004; Linder 2014). Several taxonomic lineages, including members of Pteridaceae and Polypodiaceae, exhibit notable radiations within Africa, indicating lineage-specific diversification processes across different ecological zones (Aldasoro *et al*. 2004; Smith *et al*. 2006), whereas other groups are characterized by scattered relicts and limited diversification (e.g., Thelypteridaceae, Fawcett and Smith 2021). These lineages also differ in their evolutionary branch lengths, where some groups/clades show long branches representing ancient, persistent lineages with limited recent diversification, while others display shorter branches indicative of more recent and rapid speciation events (Pryer *et al*. 2004; Schuettpelz and Pryer 2007). Such variation contributes to differences in phylogenetic alpha diversity across communities, as assemblages dominated by distantly related taxa tend to exhibit higher phylogenetic diversity than those composed of closely related species (Faith 1992; Webb *et al*. 2002; Pryer *et al*. 2004; Graham and Fine 2008). At broader spatial scales, phylogenetic beta diversity captures the turnover of evolutionary lineages among regions and provides insights into the ecological and evolutionary processes structuring biogeographic patterns (Graham and Fine 2008; Anderson *et al*. 2011). In Africa, strong environmental gradients between humid forests, montane systems, and arid lowlands are likely to drive pronounced lineage turnover, resulting in distinct phylogenetic assemblages across regions. These patterns may reflect the combined influence of environmental filtering, dispersal limitation, and historical isolation, yet they remain insufficiently quantified at the continental scale (Guevara Andino *et al*. 2021).

Despite an increasing number of studies on African ferns, most studies have focused on smaller lineages and/or limited geographic regions, with diversity relatively well-documented in certain regions, and poorly known in others. In particular, there is still limited synthesis linking species distribution with phylogenetic structure, especially in relation to both phylogenetic alpha and beta diversity across environmental gradients and historical contexts (Graham and Fine 2008). Furthermore, the roles of climatic fluctuations, refugial stability, and evolutionary processes in shaping continental-scale fern diversity patterns have not been fully integrated into a coherent framework for Africa (Linder 2014). This gap limits a comprehensive understanding of how ecological and evolutionary processes jointly structure fern assemblages across the continent. The present study investigates phylodiversity patterns of ferns in Africa, providing the most updated study from both long-standing and recently generated occurrence datasets. We identify and map the centers of taxonomic and phylogenetic alpha and beta of African ferns and investigate relationships with contemporary environmental conditions, and discuss their implications for African biogeography. This will help broaden scientific understanding of ecological and evolutionary processes that shape current fern diversity in an ancient continent.

## Methods

### Study area

As defined by the World Geographical Scheme for Recording Plant Distributions (WGSRPD), our study area included the mainland Africa as well as adjacent oceanic islands and archipelagos including Canary Islands, Madagascar, Mayotte, Mauritius, Reunion, Madeira, St. Helena, Ascension, and Tristan de Cunha archipelago, Seychelles, Comoros, Bazaruto, Zanzibar, Cape Verde, Equatorial Guinea, and São Tomé and Príncipe, as well as subantarctic islands of Prince Edwards and Marion. This region encompasses a wide range of the major physiographic and ecological zones, including the tropical rainforest of Central and West Africa, montane systems such as the Ethiopian Highlands and East African Mountains, savanna and woodland ecosystems, Mediterranean-type regions in northern and southern Africa, and extensive arid and semi-arid zones such as the Sahara and Sahel. These diverse physiographic regions reflect strong gradients in climate, elevation, and habitat heterogeneity, which may play a key role in shaping patterns of species richness and phylogenetic diversity across the continent.

### Species data

We downloaded occurrence records for African ferns from the Global Biodiversity Information Facility (GBIF; *n*=253,882 records), Symbiota African Plants Portal (https://african-plants.org/portal/index.php; *n*=21,770 records), African Plants Database (*n*=44 records), RAINBIO (*n*=16,180 records; Gilles *et al*. 2016), and recently digitized specimens across 31 U.S. herbaria (*n*=6,538 records), resulting in 298,414 records, representing 2,618 species (Table S1). To identify misidentifications, we standardized species names to the Catalog of Life (Bánki *et al*. 2026), which includes country-level distribution data for each species based on the World Ferns database (Hassler 2026), and removed any species with an exclusively non-African distribution. We cleaned the remaining records using CoordinateCleaner R package version 3.0.1 (Zizka *et al*. 2019) to remove duplicate records, records located in botanical gardens and museums, records at country centroids or capitals, records in the ocean, records outside of Africa, and other inaccurate geographical records. All taxonomic name resolution was performed with the taxastand R package (https://github.com/joelnitta/taxastand), which allows use of a custom taxonomic reference and fuzzy matching while accounting for orthographic variation in author names. All non-matching and fuzzily matched names were manually inspected, and manual matches specified as necessary. For the final dataset, we standardized species names using the Pteridophyte Phylogeny Group taxonomy v0.0.0.9005, which is the same as used by the phylogeny (see below). Infraspecific taxa such as varieties and subspecies were aggregated to species level. This resulted in total occurrence records of 127,980 points representing 1,095 African fern species.

### Phylogeny

We accessed and pruned the Fern Tree of Life (FTOL; Nitta *et al*. 2022) v.1.8.0 to our species occurrence dataset. FTOL includes a chronogram containing the most updated and largest representation of global ferns (*n*=6,216 species); it was built using publicly accessible plastid DNA, with species names standardized using the Pteridophyte Phylogeny Group taxonomy v0.0.0.9005. Pruning our species occurrence records to species in the tree resulted in 113,460 points representing 631 African ferns.

### Statistical analysis

To analyze patterns of biodiversity, we aggregated occurrence records into square grid cells. The choice of grain size used for aggregation is not straightforward: smaller grains more accurately capture specific environmental conditions but may be under-sampled, while larger grains result in a coarser characterization of environment but are better sampled (Thornhill *et al*. 2017; Nitta *et al*. 2022). We evaluated sampling completeness using redundancy (1 – species richness / abundance; Garcillán *et al*. 2003) across spatial grains from 10 km to 200 km, where grain size indicates the side length of each grid cell. At each grain size, we projected occurrence records into grid cells using the Africa Albers Equal Area Conic projection (ESRI: 102022) geographical coordinate system. We selected 100 km as the grain size value that maximized sampling completeness while minimizing environmental heterogeneity in our dataset (Supplemental Information). We constructed an incidence (presence-absence) community matrix of species occurrence records associated with the grid cells, resulting in 1,525 occupied cells of the 7,912 total grid cells. We computed species richness (SR) as the number of unique species in each grid cell, and weighted endemism (WE) as the relative endemism, calculated as species richness per grid cell weighted by their relative frequencies (Crisp *et al*. 2001; Linder 2001). Additionally, we calculated SR and WE for the larger species datasets containing 127,980 points for 1,095 species for spatial comparison. Phylogenetic diversity (PD) was computed as the sum of branch lengths of all species in each grid cell on the spanning path to the root of the phylogeny (Faith 1992). Phylogenetic endemism (PE) for each grid was calculated as the sum of the branch lengths divided by the inverse of the total number of grid cells where each tip and corresponding branches occur (Rosauer *et al*. 2009), while the relative phylogenetic diversity (RPD) was estimated as the ratio of observed PD to the PD on a null tree with the same topology but equal branch lengths (Mishler *et al*. 2014). Likewise, the relative phylogenetic endemism (RPE) is calculated as the PE on the original tree divided by PE measured on a null tree with equal branch lengths (Mishler *et al*. 2014; Borokini and Mishler 2026).

We conducted a two-tail randomization test to obtain statistical significance of the alpha phylodiversity measures, using the curveball randomization algorithm (1,000 replications of 10,000 iterations; Strona et al., 2014). The randomization algorithm randomly reassigns species occurrences across the occupied grid cells to create null floristic assemblies in each grid cell while maintaining the original richness of each grid cell (Strona *et al*. 2014). Grid cells with phylodiversity values that fall within the upper 2.5% of its corresponding random distribution were categorized as significantly high (p>0.975), while grid cells with values below the lower 2.5% were classified as significantly low (p<0.025). However, significance tests of PE statistically favor tips with longer branches because it confounds branch length with range size and has been gradually replaced by the categorical analysis of neo- and paleo-endemism (CANAPE, Mishler *et al*. 2014). To calculate CANAPE, we first identified grid cells with significant PE, then used RPE to separate them into one of three categories of range-restricted lineages that have either: (a) significantly low RPE, representing neo-endemism (b) significantly high RPE, representing paleo-endemism, (c) significantly high PE but non-significant RPE, representing mixed endemism (Mishler *et al*. 2014; Nitta *et al*. 2023; Lizardo and Ruggiero 2026). One shortcoming of CANAPE is that it does not indicate which branches on the tree drive patterns of phylogenetic endemism; this is particularly problematic in cases of mixed endemism, since different parts of the tree may have divergent signals. The recently developed Range-weighted Branch Length Difference (RWiBaLD) metric addresses this (Mishler *et al*. 2026). We calculated RWiBaLD scores for each branch of the tree by subtracting the branch length on the range-weighted comparison tree from the range-weighted observed tree, then classified them as neo-, paleo-, or meso-endemic following the procedure described in Mishler *et al*. (2026). Alpha phylogenetic diversity measures were calculated using canaper version 1.0.1 (Nitta *et al*. 2023) packages in R statistical software v4.5.3 (R Core Team 2026).

We calculated both taxonomic and phylogenetic beta diversity to investigate floristic dissimilarities between grid assemblies, and shared or unshared branches between locations, respectively (Graham and Fine 2008; Laffan *et al*. 2016; Borokini and Mishler 2026). In this study, we used the Sorensen index and PhyloβSørensen method to calculate taxonomic and phylogenetic beta diversity respectively, for the African ferns. Optimal *k* clusters were determined using the elbow method based on hierarchical clustering and were used for plotting maps of beta diversity patterns for bioregionalization. Beta diversity measures were computed in phylospatial R package version 1.3.0.9 (Kling 2025). Since real bioregions are not discrete, we also used the Grade of Membership model (White *et al*. 2019) to assign grid cells to bioregions with partial membership; this method also allows us to identify the taxa that are mostly strongly associated with each bioregion.

We modeled relationships among the phylodiversity measures with selected edaphic and bioclimatic variables. We downloaded monthly mean temperature (bio1), temperature seasonality (bio4), maximum temperature of the warmest month (bio5), annual precipitation (bio12), and precipitation seasonality (bio15) from the WorldClim 2.0 database (1970-2000; Fick and Hijmans 2017). These selected climatic datasets represent mean and variability conditions in the study area. We also accessed five global soil variables – soil pH, fine organic carbon, soil nitrogen, organic carbon density, and bulk density – at 5 cm depth using the soil_world function in geodata R package (Hijmans *et al*. 2026). The data is based on the soilGRIDS database and downloaded at 1 km spatial resolution. We removed any variables with Pearson’s correlation coefficient >0.7. The selected climatic and edaphic variables were resampled using the average for each grid cell to the same spatial resolution used for the phylodiversity analyses (100 km) prior to modeling. Spatial coordinates (longitude and latitude) were included as predictors to account for spatial structure, and residual spatial autocorrelation was assessed using Moran’s I statistic. For alpha diversity metrics, relationships with environmental predictors were analyzed using spatial generalized linear models implemented in sdmTMB R package version 1.0.0 (Anderson *et al*. 2025) and spatial random effects modeled using a Gaussian Markov random field with a mesh cutoff distance of 200 km. SES values of PD, PE, RPD, and RPE (standardized effect sizes relative to the null model) were modeled with a Gaussian error distribution, whereas species richness was modeled with a negative binomial distribution.

All code used in the analysis is available at https://github.com/joelnitta/africa_ferns/.

## Results

### Alpha diversity

Observed species richness for the species represented on the phylogeny showed relatively low distributions (<50 species) across most of Africa, while exceptionally high richness (≥200 species) was observed in Madagascar, and a few grid cells with moderately high species richness (>100 species) observed in the Congo Basin rainforest, eastern part of Southern Africa, the Upper Guinean Forest, and parts of the eastern arc mountains in Kenya and Tanzania (Figure 1a). Fern richness for the unfiltered dataset shows similar pattern with grid cells in Madagascar over 300 species (Figure 1b). Furthermore, the pattern of spatial phylogenetic diversity (PD) pattern (Figure 1c) is similar to the fern species richness, showing mostly low PD distributed across Africa, and scattered grid cells with relatively high PD (0.3-0.5) observed in northern and eastern Madagascar, the eastern part of Southern Africa, the Congo Basin, eastern arc mountain region, and the Upper Guinea forests (Figure 1c). The PD of African ferns was characterized by low significance (P<0.01<0.025) across the entire continent and adjacent oceanic islands, while a few cells with high PD significance (P>0.975>0.99) were scattered in the Congo Basin and the Mediterranean tips of northern and southern Africa (Figure 1d). As expected, PD and species richness of African ferns were significantly linearly related (Pearson’s correlation coefficient; r=0.98, p<0.001).

**Figure 1:**
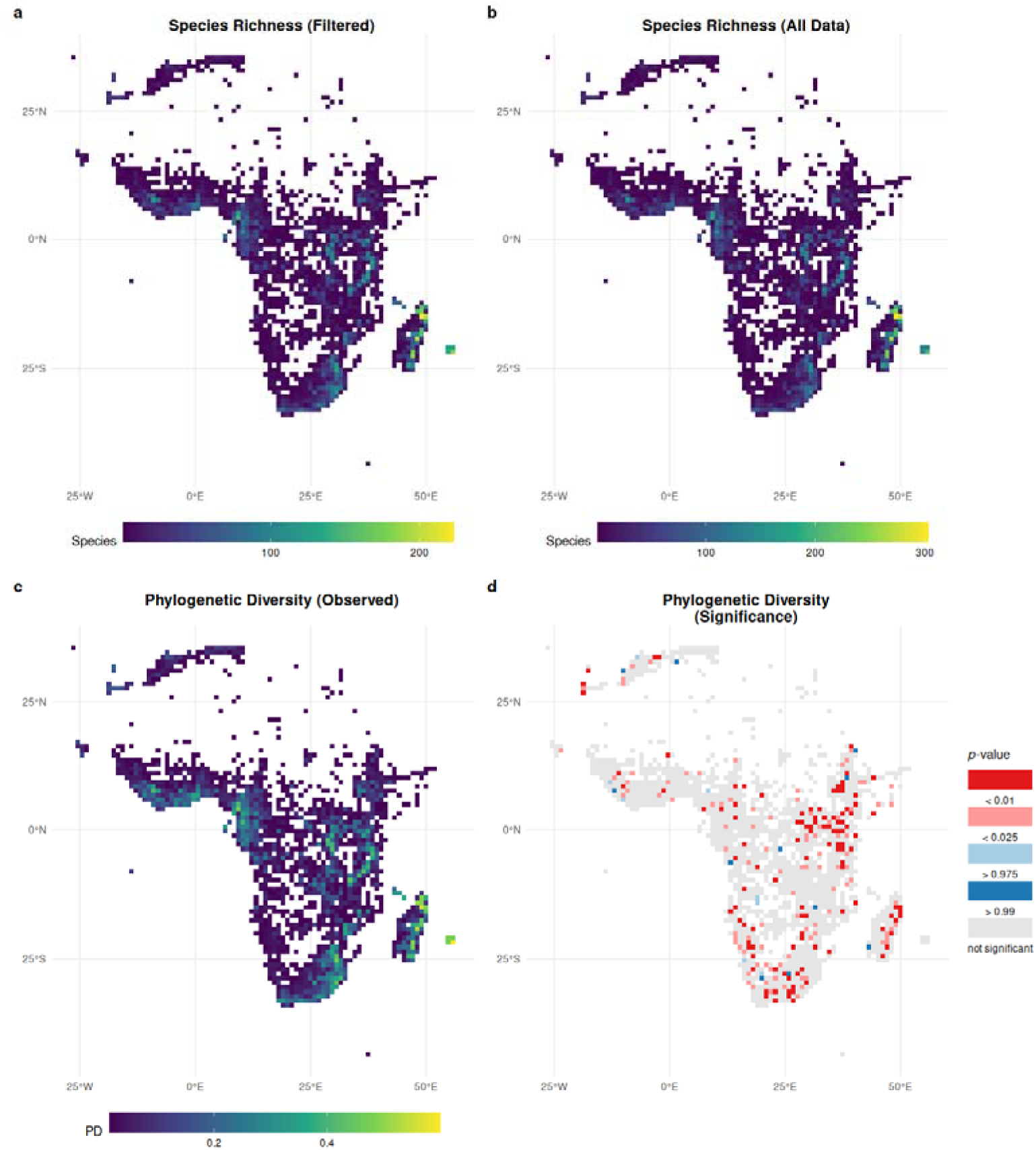
Maps of spatial patterns of alpha diversity of African ferns, representing (a) species richness for the 113,460 occurrence points (n=631 species found on the phylogeny), (b) species richness of 127,980 points representing 1,095 ferns, (c) observed phylogenetic diversity, and (d) significant phylogenetic diversity.

Weighted endemism (WE) of ferns is generally low across Africa, with relatively high WE observed only in Madagascar (Fig. S3). Like the WE, phylogenetic endemism (PE) was low across most of Africa, however, grid cells with high PE were observed in Madagascar, the Mediterranean North Africa, and adjacent oceanic islands (Figure 2a). Low PE significance (P<0.01<0.025) was observed across the Guinean and Congolian tropical rainforest ecosystems, the eastern arc mountains, and the Ethiopian highlands (Figure 2b). Raw PE and WE patterns were also highly positively correlated (Pearson’s correlation coefficient; r=0.99, p<0.001).

**Figure 2:**
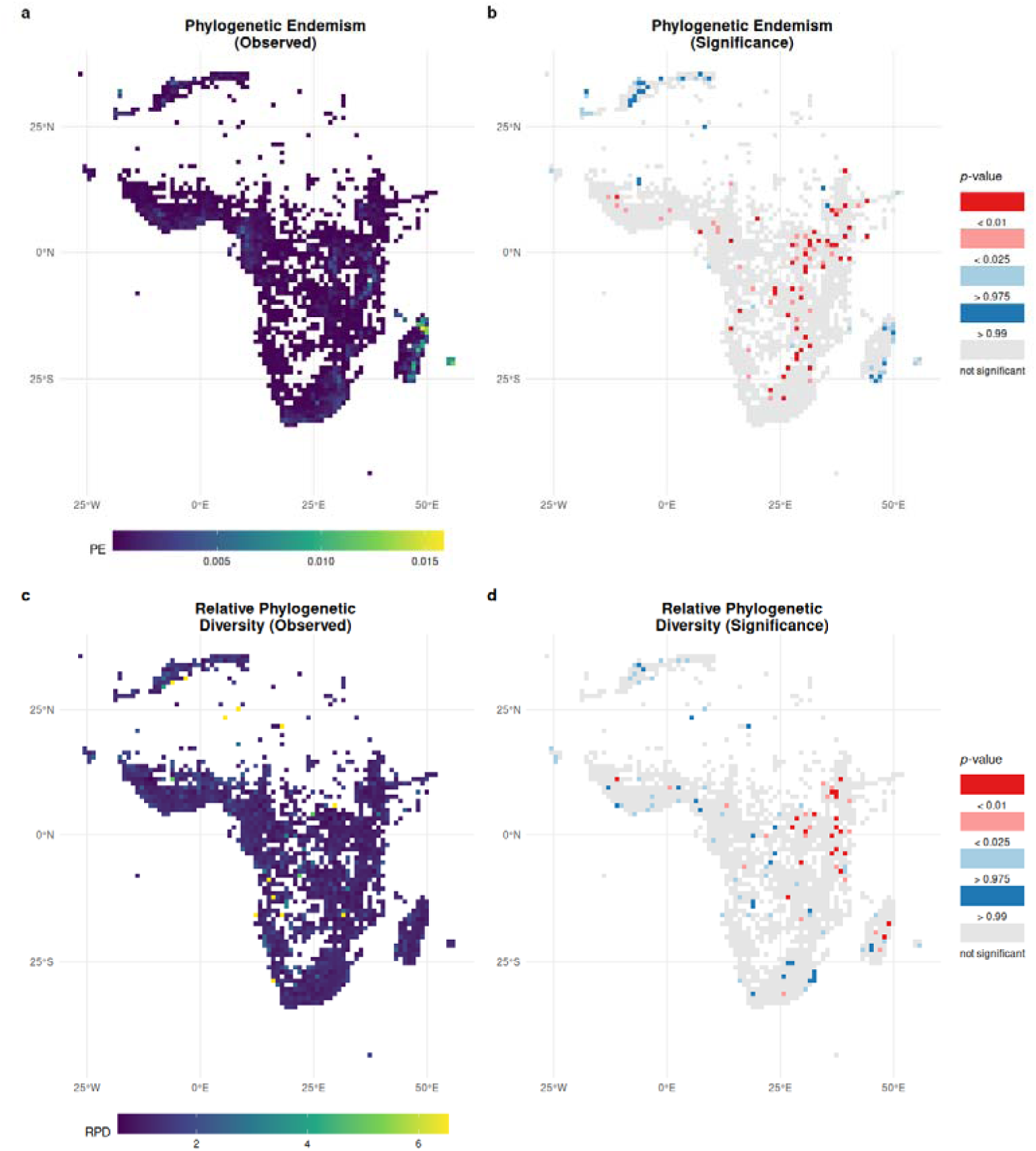
Maps of spatial patterns of phylogenetic endemism and relative diversity of African ferns (n=113,460 occurrence points representing 631 species found on the phylogeny). (a) Observed phylogenetic endemism, (b) statistical significance of phylogenetic endemism, (c) observed relative phylogenetic diversity, and (d) significance of relative phylogenetic diversity.

Spatial patterns of relative phylogenetic diversity (RPD) of African ferns were characterized by low values across the continent while the highest RPD values were scattered in the Sahara Desert, lower Congo Basin of Angola, and the Mediterranean North Africa (Figure 2c). Grid cells with intermediate RPD values (3-4) were noted across semi-arid areas of the Guinean forests and arid regions of southern Africa as well as the Mediterranean North Africa (Figure 2c). Grid cells of low RPD significance (P<0.01<0.025) were distributed in the savanna ecosystems of central Africa, the Ethiopian Highlands, eastern arc mountains, and eastern Madagascar, while grid cells of high RPD significance (P>0.975>0.99) were concentrated in the arid and semi-arid regions of Southern Africa, western Madagascar, Guinean forests of West Africa, savanna ecosystems of south-central Africa, and the Mediterranean North Africa (Figure 2d).

The most frequent type of endemism in African ferns identified by CANAPE analysis was mixed endemism (co-occurrence of phylogenetically young and older lineages, or in some cases lineages of intermediate age; green cells) primarily concentrated in Mediterranean North Africa and Madagascar, as well as oceanic islands of Mayotte, Canary Islands, and Cape Verde, and were associated with a range of biomes (Figure 2). Neo-endemic cells were primarily concentrated in Madagascar, with additional occurrences in the lower Congo Basin and along the banks of the Kunene River (Figure 2). The few paleo-endemic cells were scattered in the Guinean forests and Congolian Basin (Figure 2).

Range-weighted branch length difference (RWiBaLD) analysis revealed that cells of mixed endemism identified by CANAPE comprised a mixture of intermediate (meso) and young (neo) branches (Fig. S4). However, within tropical moist forests on Madagascar, some cells that were identified by CANAPE as mixed included a preponderance of neo-endemic lineages, while within Mediterranean Morocco and Madeira some mixed cells included most paleo-endemic lineages (Figure 4). Neo-endemic lineages were mostly represented in tropical moist forests, while the paleo-endemic lineages were distributed across the Mediterranean, desert, and tropical moist forest biomes (Figure 3). When plotting RWiBaLD endemism status on the tree (Fig. S5), a total of 23 older and range-restricted (that is, paleo-endemic) terminals were observed, including *Asplenium hispanicum*, *A. fontanum*, *Gymnocarpium robertianum*, *Notogrammatis angustifolia*, and *Myriopteris microphylla*, among others. However, there were more phylogenetically young and range-restricted terminals on the tree (*n*=97 tips), while 58 meso-endemic tips and 7 internal branches were observed, but the internal meso-endemic branches diversified into neo-endemic fern tips (Fig. S5).

**Figure 3:**
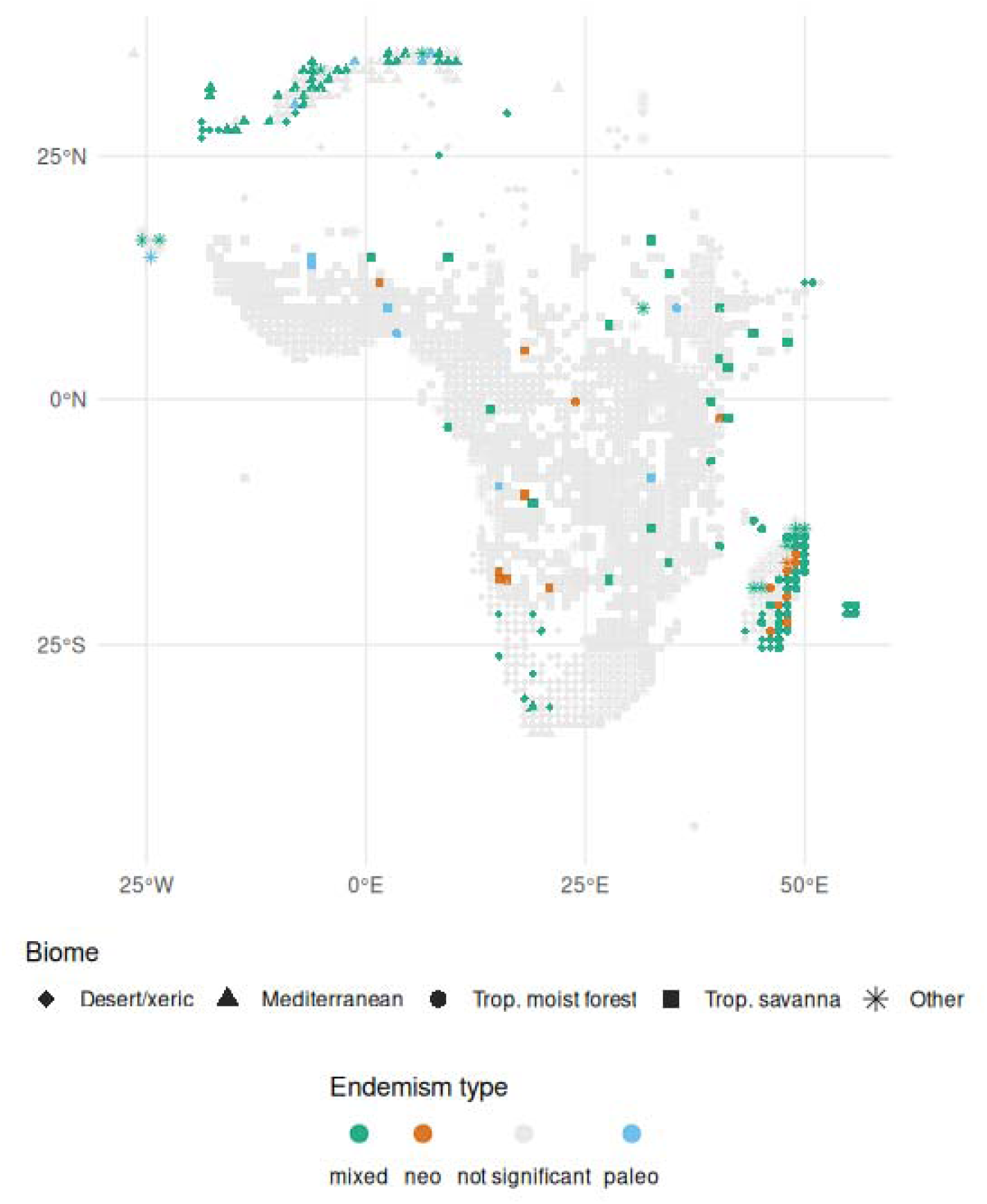
CANAPE endemism classification for African fern grid cells, with each cell shown as a point colored by endemism type and shaped by biome (rare biomes grouped as Other).

**Figure 4:**
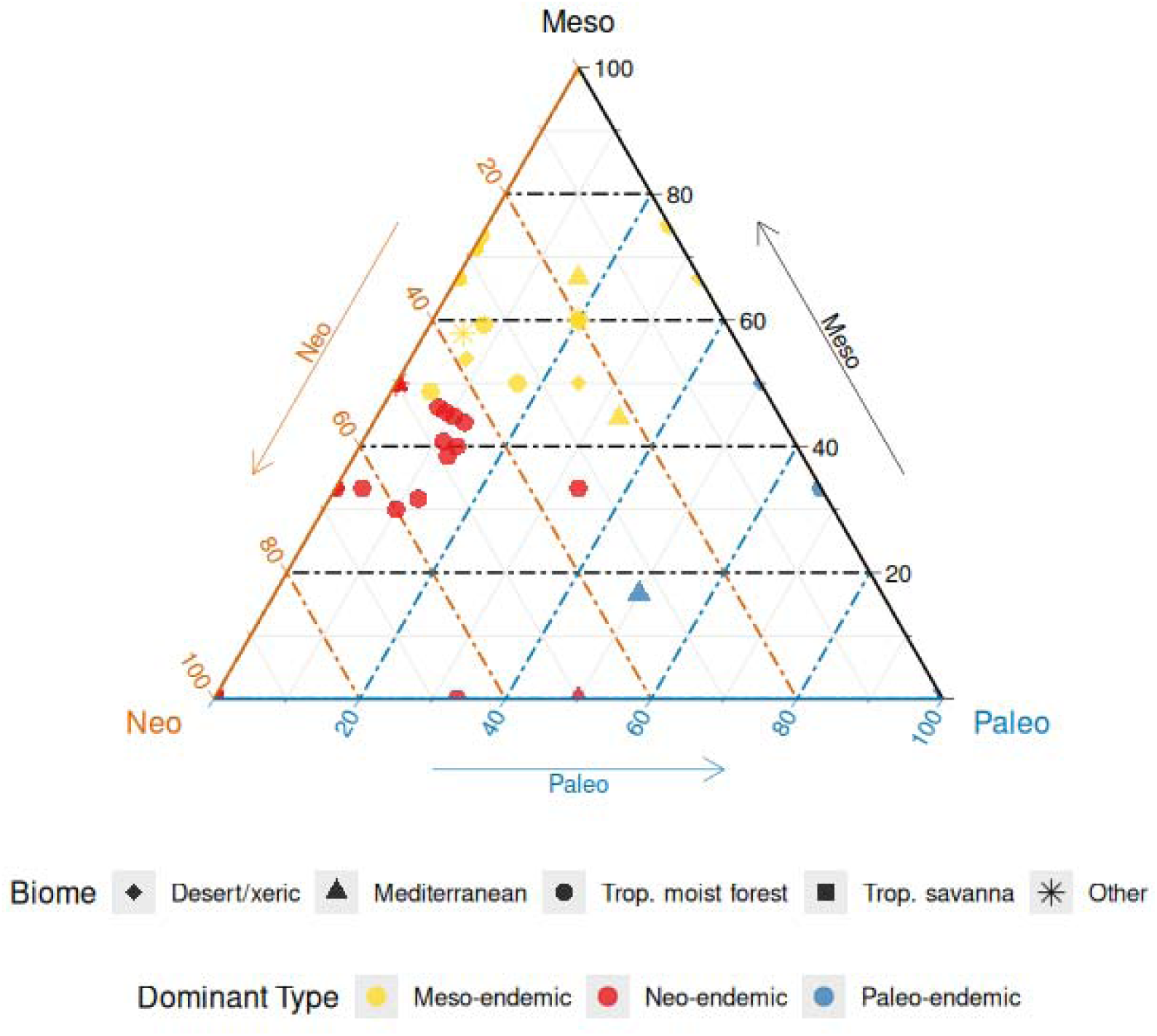
Ternary plot showing the endemism composition of mixed endemism grid cells as determined by range-weighted branch length difference (RWiBaLD) analysis. Each axis shows the percentage of key branches classified as neo-endemic (Neo), meso-endemic (Meso), or paleo-endemic (Paleo); unclassified key branches are excluded from both the percentages and the totals. Each point is one grid cell; color indicates the dominant endemism type and shape indicates biome.

### Beta diversity and bioregionalization

The regionalization of taxonomic and phylogenetic beta diversity (Figure 5a–b, respectively) of African ferns using hierarchical clustering produced vastly different results. Taxonomic bioregion 1 (black) was concentrated in Mediterranean North Africa, adjacent Canary Islands, semi-arid western Africa and central Africa, southern Africa, along the East Africa coastline, and eastern Madagascar. Taxonomic bioregion 2 (orange) was mainly distributed in eastern Africa, Ethiopian highlands, savanna ecosystem of Central Africa, most of the Cape Floristic Province, and the western half of Madagascar, the adjacent Mayotte, Reunion, Mauritius, and lower Congo Basin. Taxonomic bioregion 3 (blue) dominated the arid southern Africa (Kalahari and Namib Deserts) and the Horn of Africa, Cape Verde, and a few cells in eastern Madagascar, while taxonomic bioregion 4 (green) was primarily distributed in the Guinean Forests of West Africa and the Congo Basin (Figure 5a).

**Figure 5:**
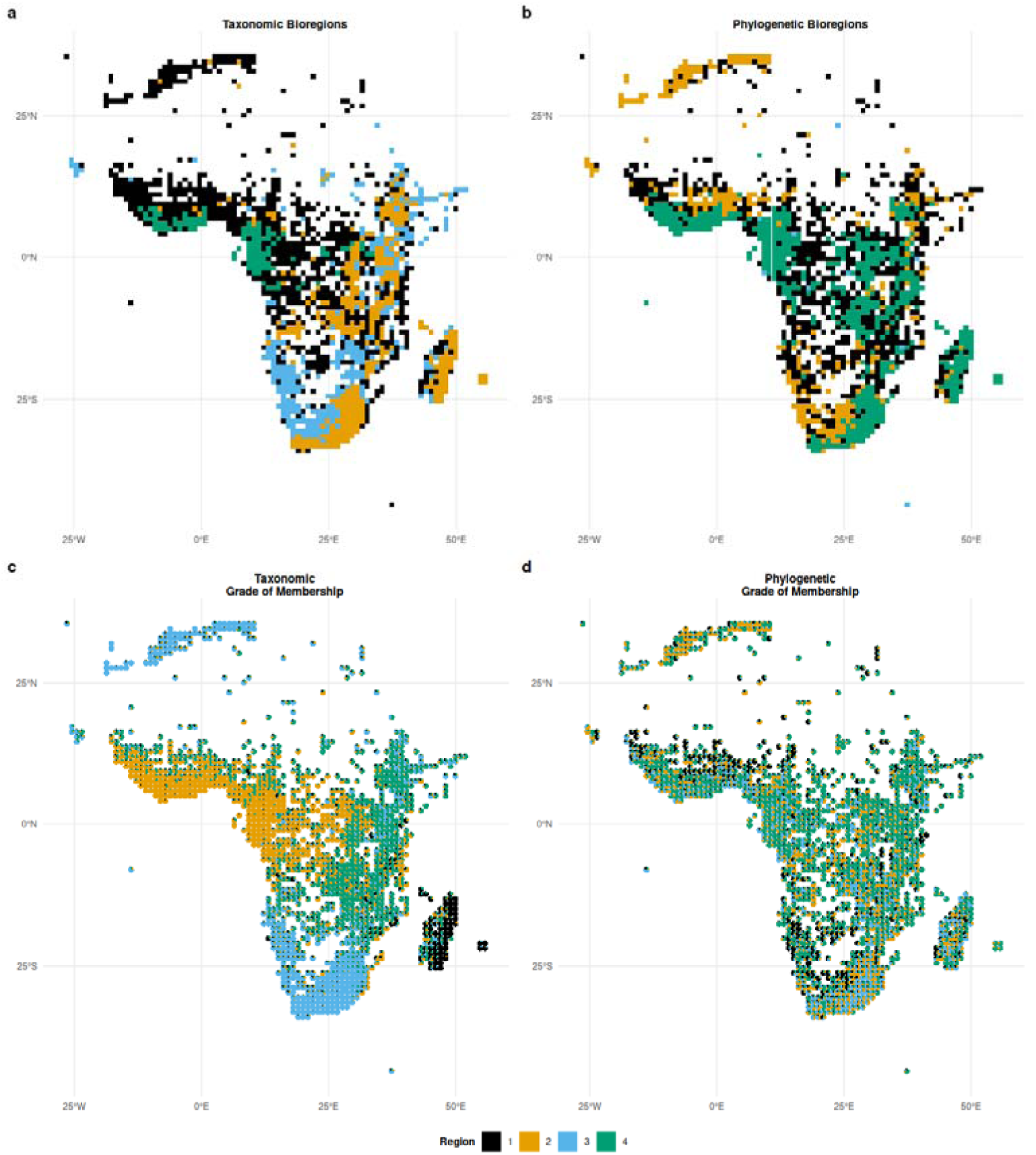
Bioregionalization of African ferns (n=113,460 occurrence points for 631 species). (a) Taxonomic and (b) phylogenetic beta diversity regionalization based on hierarchical clustering. (c) Taxonomic and (d) phylogenetic Grade of Membership (GoM) regionalization. Note that although a single color scheme is used for all plots, each bioregionalization is independent.

Phylogenetic regionalization showed that region 1 (black) has strong affinity with semi-arid West Africa and parts of the Kalahari and Namib Deserts, parts of the Congo Basin, western Madagascar, and lower eastern Africa, while region 2 (orange) is concentrated in Mediterranean North Africa and adjacent oceanic islands, semi-arid West Africa, and the desert regions of southern Africa (Figure 5b). Region 3 (blue) is represented by fewer than 10 grid cells found in St. Helena, Ascension, and Tristan de Cunha archipelago, Equatorial Guinea, as well as central and eastern Africa, while region 4 (green) is the most widely distributed phyloregion cluster across the continent south of the Sahara Desert (Figure 5b).

The grade of membership (GoM) model also recovered more discrete regionalization for taxonomic vs. phylogenetic distances, but the spatial distribution differed from the hierarchical clustering results. GoM taxonomic bioregion 2 roughly corresponded to hierarchical taxonomic bioregion 4, but South Africa and Mediterranean North Africa were both associated with GoM taxonomic bioregion 3, much of the eastern part of the continent was associated with GoM taxonomic bioregion 4, and GoM taxonomic bioregion 1 was largely confined to Madagascar (Figure 5c). The GoM model based on phylogenetic distances showed very little spatial signal, with a near even mixing of bioregions across the study area (Figure 5d).

### Drivers of phylodiversity patterns

Results of the spatial generalized linear models including geographical distance as a random effect are shown in Figure 6 and Fig. S6. PD SES decreased significantly along the gradients of soil pH and soil nitrogen, but other variables had no significant effect; however, none of the variables had a significant effect on PE SES (Figure 6). Furthermore, RPD SES increased with temperature seasonality and annual mean temperature, but decreased with maximum temperature of the warmest month. In contrast, RPE SES was significantly driven only by soil pH. Finally, species richness decreased with maximum temperature of the warmest month and soil bulk density (Figure 6).

**Figure 6:**
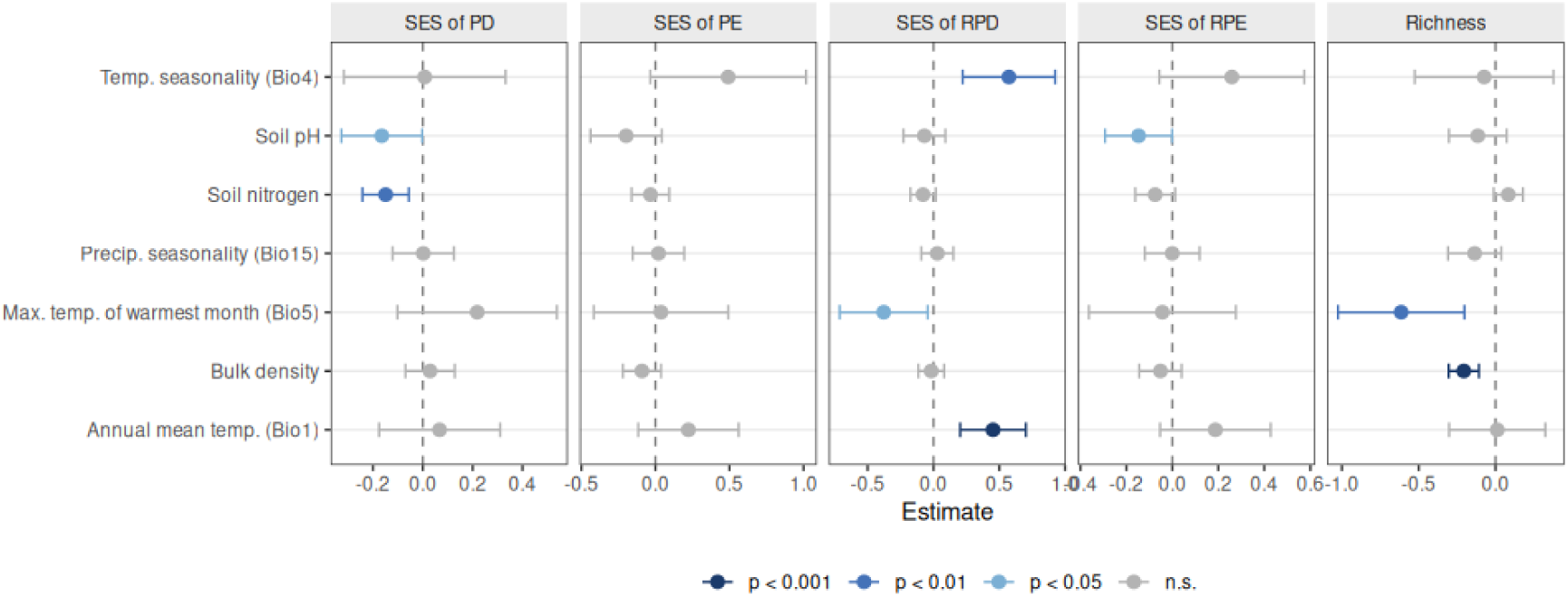
Relationship of alpha diversity metrics with environmental variables. Points show model estimates ± 95% CI; colours indicate significance level.

## Discussion

We mapped the centers of species and phylogenetic diversity and endemism of fern flora in Africa. Fern richness and diversity on the continent is generally low but also disproportionately distributed, with peak richness and diversity observed in Madagascar and the adjacent oceanic islands. Our finding supports previous studies that have reported low fern richness in Africa (e.g., Tryon 1986; Kornaś 1993; Aldasoro *et al*. 2004). Furthermore, fern phylogenetic diversity was broadly characterized by significant concentrations of closely related taxa (that is, phylogenetic clustering), while significantly high species and phylogenetic endemism were observed mainly in Mediterranean North Africa and Madagascar and adjacent Indian Ocean islands. Older fern lineages were scattered across the continent and adjacent islands, while younger lineages were concentrated in the alpine areas of the Eastern Arc Mountains and the Ethiopian Highlands, and eastern Madagascar, suggesting that these regions are evolutionary cradles of fern speciation. Centers of phylogenetic endemism were primarily concentrated in Madagascar, eastern Africa, and northern Africa, and were characterized by both neo- and paleo-endemic fern lineages (that is, mixed endemism). The grid cells of mixed endemism were characterized by both ferns with phylogenetically intermediate and young ages. These spatial phylogenetic patterns are significantly influenced by both climatic and edaphic factors.

This study addresses a key gap in understanding how species distributions and evolutionary history jointly shape African fern diversity. By combining taxonomic and phylogenetic perspectives, we show that fern diversity across the continent reflects a complex interplay between environmental gradients, historical isolation, deep biogeographical events, and lineage-specific diversification. The observed mismatch between species richness and phylogenetic diversity shows that regions of high richness do not necessarily represent centers of evolutionary distinctiveness, emphasizing the importance of integrating α- and β-diversity to interpret large-scale biodiversity patterns.

### Spatial and evolutionary patterns of African fern diversity

The spatial patterns of both the species and phylogenetic richness of the African ferns show relatively high diversity in the tropical and subtropical biomes, island systems, and areas with microclimates driven by large adjacent aquatic habitats. Spatial patterns of fern diversity across Africa reflects strong regional controls driven by the combined influence of moisture availability, topographic complexity, habitat area, and biogeographic history (Mehltreter *et al*. 2010; Karichu *et al*. 2022, 2024). For instance, in Madagascar, high fern species richness is concentrated in the humid eastern and northeastern regions, including Montagne d’Ambre National Park, Marojejy National Park, Ambohitantely National Reserve, the Andasibe–Maromizaha corridor, and Vohimana, where persistent rainfall, steep elevational gradients, and structurally complex montane and lowland forests create a wide range of microhabitats that support fern establishment and persistence (Rakotondrainibe 2000; Randrianarimanana *et al*. 2024). The relatively large extent and continuity of humid forest in these regions, coupled with long-term climatic stability, have further allowed the accumulation and persistence of both widespread and localized species, reinforcing their role as diversity centers (Janssen *et al*. 2007; Kessler *et al*. 2011).

Fern richness in Mainland Africa is concentrated in montane systems such as the Eastern Arc Mountains and Ethiopian Highlands, where high and persistent moisture, steep elevational gradients, and habitat heterogeneity generate a wide range of ecological niches that support species accumulation (Hemp 2002; Liu *et al*. 2016; Suissa *et al*. 2021). In West Africa, fern richness is largely concentrated in the Lower Guinean forests and the Cameroon Highlands, which function as long-standing humid refugia (Maley 1996), where high rainfall, elevational gradients, and habitat heterogeneity support species accumulation, with over 100 species recorded on Cameroon Mountain alone (Tryon 1965). Central African richness patterns are largely driven by the Congo Basin, where extensive lowland rainforest provides large, continuous areas of suitable habitat and stable microclimatic conditions for fern persistence (Kornaś *et al*. 2000). In southern Africa, relatively higher fern richness in the Cape region reflects localized moisture availability and topographic complexity (Aldasoro *et al*. 2004), which creates refugial conditions despite the broader arid context.

Arid and semi-arid regions of Africa, characterized by low precipitation, high temperatures, and strong seasonality, can only support low fern richness due to limited gametophyte survival and reproductive success (Page 2002; Kessler *et al*. 2011), as reflected in our results. Where present, such as in North Africa, ferns are largely confined to localized abiotic refuges and higher elevations that buffer environmental stress (Anthelme *et al*. 2011).

The spatial patterns of relative phylogenetic diversity (RPD) significance show a concentration of younger fern lineages in the Eastern Arc Mountains, the Ethiopian Highlands and adjacent areas, indicating that these areas are centers of recent speciation and cradles of diversification of African ferns, while significant concentration of older lineages across the continent highlight refugia hotspots (Figure 2d). The Eastern Arc Mountains harbor over 800 species of endemic vascular flora (Burgess *et al*. 2007) and has been noted as a “cradle of diversity” but also as a refugia and diversification museum in previous studies (Roy 1997; Dimitrov *et al*. 2012; Dagallier *et al*. 2020). While the Eastern Arc Mountains resulted from the Precambrian uplift over three billion years ago, some of the areas have been covered by volcanic and sedimentary rocks from recent (<5 million years BP) geological events (Newmark 2002; Burgess *et al*. 2007). The Ethiopian Highlands share a similar geological history of being formed from Precambrian volcanoes but overlaid by more recent Cenozoic uplifts and volcanic basaltic lava (Merla *et al*. 1979; Lanckriet *et al*. 2015). These recent geological events created novel habitats that may have shaped the flora of the region, resulting in more recent diversification of fern lineages, many of which are endemic. In contrast to mountainous regions, large areas of low-elevation arid habitat that may have supported more ferns under previous climates are now inhospitable to most species. Repeated shifts in climate likely drove many lineages to extinction and restricted others to refugia (Aldasoro *et al*. 2004). These events have led to the evolution of unique African lineages, e.g., *Menisorus* (Thelypteridaceae; Fawcett and Smith 2021) and the African *Platycerium* radiation (Polypodiaceae; Janssen *et al*. 2007), while also providing ecological opportunity for ancient and recent colonization from Asia and the Americas (Moran and Smith 2001; Janssen *et al*. 2007).

### Patterns of Endemism

Endemism in African ferns is strongly spatially structured, with both species weighted and phylogenetic endemism identifying localized hotspots in the Canary Islands, the Lower Guinean forests of West Africa, eastern and southern Africa, and Madagascar. These regions represent centers where range-restricted taxa accumulate alongside lineages with diverse evolutionary histories, reflecting the combined influence of ecological specialization and historical isolation (Barrington 1993; Janssen *et al*. 2007; Suissa *et al*. 2021).

Across much of the continent, mixed endemism (co-occurrence of neo- and paleo-endemics) (Mishler *et al*. 2014), indicates that African fern assemblages reflect both recent diversification and the persistence of older lineages (Sessa *et al*. 2017; Loiseau *et al*. 2020; Howard *et al*. 2025), pointing to multiple evolutionary processes rather than a single dominant diversification pathway (Qian *et al*. 2023; Qian 2024). This is further supported by the spatial partitioning of neo-endemism in eastern and southern Africa and Madagascar, and paleo-endemism in northern and western regions.

However, the spatial signal of low PD significance across large parts of western, central, and southern Africa, as well as Madagascar, indicates clustering of closely related fern taxa, suggesting that endemism in these regions is largely driven by recent radiations or environmental filtering rather than deep evolutionary divergence (Crisp *et al*. 2001; Cavender-Bares *et al*. 2009). This pattern likely reflects recent radiations within specific clades, such as African Polypodiaceae (Janssen *et al*. 2007), cheilanthoid ferns in the Cape Floristic Region (Eiserhardt *et al*. 2011), and scaly tree ferns in Madagascar (Janssen *et al*. 2008), resulting in assemblages composed of relatively closely related lineages.

Mediterranean North Africa was recovered as an area of high paleo-endemism in our results (Figure 2b; Figure 3). However, unlike the highlands of Madagascar and South Africa, which are isolated from other areas of equable habitat and host many endemics, montane areas of North Africa (e.g., the Atlas Mountains) serve as the southern range limit for many species also distributed in Europe and/or the Middle East. Some of these (e.g., *Gymnocarpium robertianum*) are circumboreal and have isolated stations in the high montane areas, while others (e.g., *Asplenium hispanicum*) are endemic to the Mediterranean basin.

### Beta Diversity in African Ferns

Species-based biotic regionalization of African ferns computed using the grade of membership model is more resolved than the clustering analysis. Species-based GoM showed strong climatic and geographic structuring of the regions. Region 1 (black) is restricted to the island nations of the Indian Ocean, region 2 (orange) is associated with the Guineo-Congolian Afrotropical rainforest ecosystems, while regions 3 (blue) and 4 (green) are restricted to the northern and southern Africa deserts and tropical grassland biome, respectively (Figure 5c). Additionally, all four regions, except region 3, exhibit high geographic structure, while the spatial distribution of all species-based GoM regions are loosely congruent with Africa’s vegetation zones (Fig. S2). The spatial patterns in compositional turnover indicate significant climatic underpinning in the distribution of African ferns, highlighting the role of environmental preference and niche sorting in African fern assemblages (Webb *et al*. 2002; Mienna *et al*. 2020). Ecological affinities are clearly pronounced in West Africa where GoM for the rainforests are different from the semi-arid savanna where the latter is shared with the eastern Africa-dominated GoM. A similar pattern of ecological association of GoM regions was observed for western and eastern Madagascar.

GoM model for the phylogenetic-based regionalization of African ferns shows different patterns from the species-based GoM. Phylogenetic-based GoM for African ferns show continental distributions of regions 2-4 (orange, blue, and green, respectively), inclusive of Madagascar and adjacent oceanic islands, indicating deep biogeographic events and shared evolutionary history of ferns across the continent. Moreover, the shared phylogenetic branches between mainland Africa and Madagascar further support earlier reports of fern speciation in Madagascar and dispersal into the African continent (Bauret *et al*. 2017; Hennequin *et al*. 2017). While GoM region 4 is widely distributed across the continent, with higher membership in the tropical rainforest and savanna ecosystems, greater partial representations of GoM regions 1-3 were noted in restricted and relatively small parts of the continent. Region 1 (black) was more represented in the savanna ecosystem of northern West Africa and the desert landscapes of southern Africa, the Mediterranean northeastern Africa have higher representations of region 2 (orange), while region 3 (blue) was observed in southern Africa (Figure 5d). Overall, the phylobeta GoM of African ferns do not reflect ecological, climatic, or geographic affinities, which may be due either to the presence of widespread ferns or strong evolutionary signal associated with taxa with long branches (Scherson *et al*. 2017).

### Limitation of the study

The inferences of the study are limited partly by a lack of herbarium collection data across much of the continent, particularly interior equatorial humid-tropical regions of the Congo Basin, and consequent bias of collection density in more populous and better-explored regions. About 53% of the fern flora is represented in our phylogeny, which may have led to the omission of short-branch neoendemics. Furthermore, the taxonomic distribution dataset and the phylogenetic dataset are derived from only partly overlapping sources, so widespread species occurring in Africa are often represented by sequence data from a sample collected outside of Africa, and may be subject to misidentification or overly-broad application of species concepts. Despite these limitations, a clearer picture has emerged of the broad patterns of fern diversity and endemism, highlighting areas of high species richness and the evolutionary processes that have contributed to their assembly.

## Acknowledgments

Some of the dataset used in this study is based upon work supported by the National Science Foundation under the following DBI grant award numbers: DBI-2223875 (Kansas University), DBI-2223876 (University of California Berkeley), DBI-2223877 (Rancho Santa Ana Botanic Garden), DBI-2223878 (Arizona State University), DBI-2223879 (Missouri Botanical Garden), DBI-2223880 (Harvard University), and DBI-2223881 (New York Botanical Garden). Any opinions, findings, and conclusions or recommendations expressed in this material are those of the author(s) and do not necessarily reflect the views of the National Science Foundation.

## Supplemental Information

**Table S1:** Number of fern species per country in the full dataset (all cleaned occurrence records) and the pruned dataset (species also present in the phylogenetic tree).

| Country | Full dataset | Pruned dataset |
|---|---|---|
| Madagascar | 648 | 403 |
| Tanzania | 375 | 275 |
| Democratic Republic of the Congo | 336 | 240 |
| Cameroon | 293 | 210 |
| South Africa | 275 | 212 |
| Kenya | 272 | 220 |
| Uganda | 218 | 175 |
| Réunion | 209 | 188 |
| Equatorial Guinea | 201 | 153 |
| Ethiopia | 200 | 164 |
| Mozambique | 187 | 157 |
| Gabon | 185 | 134 |
| Burundi | 174 | 134 |
| Zimbabwe | 170 | 138 |
| Côte d'Ivoire | 157 | 128 |
| Guinea | 156 | 122 |
| Malawi | 152 | 129 |
| Rwanda | 144 | 109 |
| Ghana | 140 | 115 |
| Liberia | 138 | 115 |
| Zambia | 135 | 105 |
| Nigeria | 123 | 102 |
| Angola | 120 | 102 |
| Comoros | 116 | 108 |
| Republic of the Congo | 108 | 86 |
| São Tomé and Principe | 106 | 92 |
| Togo | 96 | 77 |
| Kingdom of eSwatini | 91 | 78 |
| Mayotte | 89 | 84 |
| Sierra Leone | 89 | 78 |
| Lesotho | 70 | 62 |
| Central African Republic | 67 | 55 |
| Namibia | 59 | 51 |
| Madeira | 53 | 48 |
| Benin | 49 | 40 |
| South Sudan | 49 | 43 |
| Morocco | 47 | 44 |
| Spain | 45 | 41 |
| Botswana | 37 | 30 |
| Algeria | 35 | 34 |
| Sudan | 32 | 29 |
| Republic of Cabo Verde | 31 | 29 |
| Senegal | 30 | 24 |
| Burkina Faso | 26 | 22 |
| Eritrea | 21 | 17 |
| Mali | 21 | 17 |
| Tunisia | 18 | 18 |
| Chad | 17 | 17 |
| Zanzibar | 17 | 17 |
| Saint Helena | 15 | 15 |
| Guinea-Bissau | 13 | 12 |
| Somalia | 13 | 11 |
| Somaliland | 12 | 9 |
| Egypt | 7 | 6 |
| Niger | 7 | 5 |
| The Gambia | 6 | 4 |
| Djibouti | 5 | 3 |
| Libya | 3 | 3 |
| Mauritania | 3 | 3 |
| Azores | 1 | 1 |

**Table S2:**
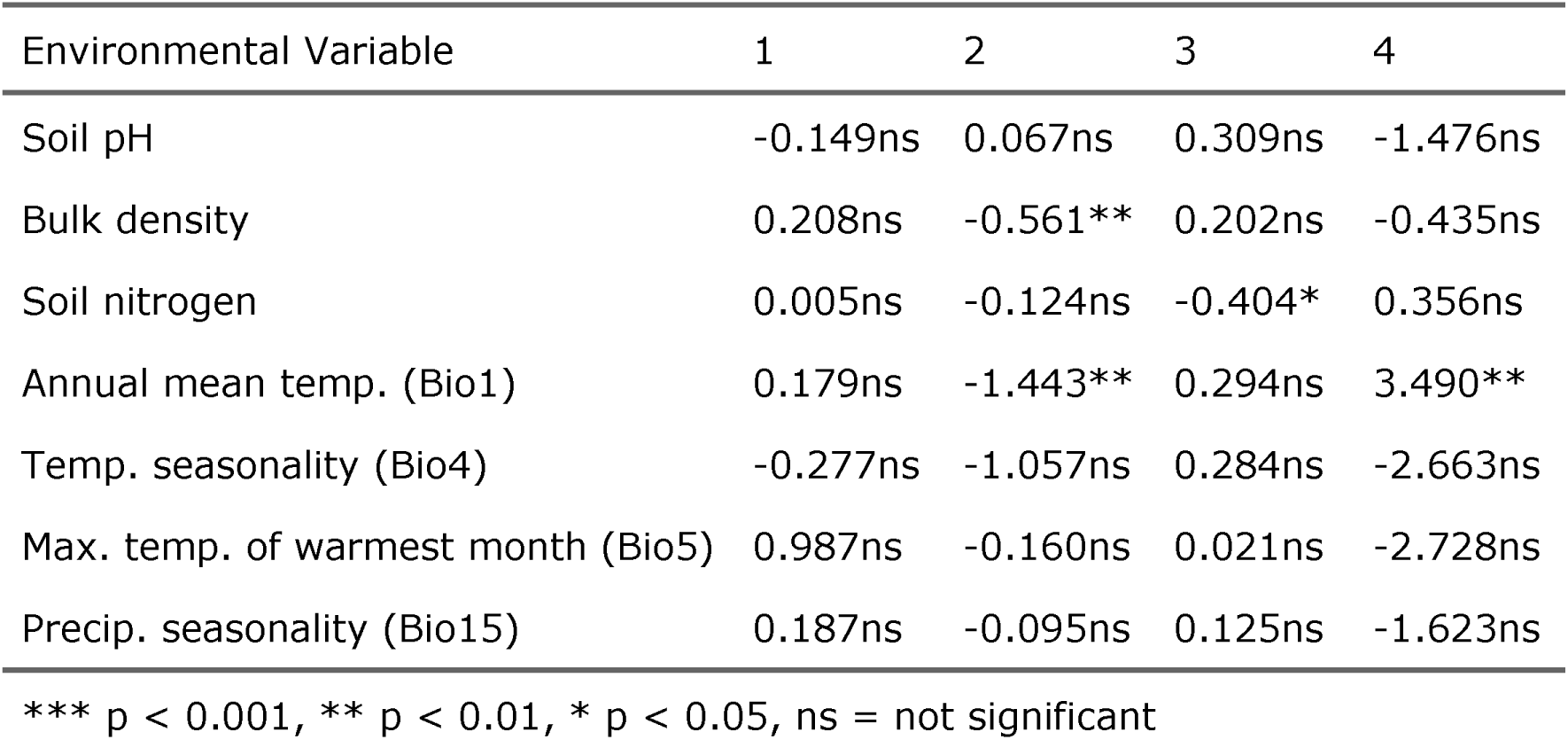
Relationship of taxonomic hierarchical clustering bioregions with environmental variables.

**Table S3:** Relationship of phylogenetic hierarchical clustering bioregions with environmental variables.

| Environmental Variable | 1 | 2 | 3 | 4 |
|---|---|---|---|---|
| Soil pH | 0.094ns | 0.645* | 0.389ns | -0.442ns |
| Bulk density | 0.255* | -0.012ns | 0.033ns | -0.348* |
| Soil nitrogen | -0.231ns | -0.090ns | -0.438ns | 0.227ns |
| Annual mean temp. (Bio1) | -0.124ns | -0.898* | 1.832ns | 0.421ns |
| Temp. seasonality (Bio4) | -0.658ns | -0.200ns | 0.582ns | -0.257ns |
| Max. temp. of warmest month (Bio5) | 1.198** | 0.509ns | -1.241ns | -1.711** |
| Precip. seasonality (Bio15) | 0.128ns | 0.249ns | -1.121* | -0.225ns |

\*\*\* p < 0.001, \*\* p < 0.01, \* p < 0.05, ns = not significant

**Table S4:** Relationship of taxonomic GoM topics with environmental variables.

| Environmental Variable | 1 | 2 | 3 | 4 |
|---|---|---|---|---|
| Soil pH | 0.011ns | -0.047** | 0.039** | -0.011ns |
| Bulk density | -0.014ns | 0.001ns | 0.013ns | 0.002ns |
| Soil nitrogen | 0.002ns | 0.003ns | -0.002ns | -0.006ns |
| Annual mean temp. (Bio1) | -0.121*** | 0.107*** | -0.020ns | 0.035* |
| Temp. seasonality (Bio4) | -0.058ns | 0.032ns | -0.065ns | 0.042ns |
| Max. temp. of warmest month (Bio5) | 0.094* | -0.067ns | 0.005ns | -0.025ns |
| Precip. seasonality (Bio15) | 0.061*** | -0.015ns | -0.012ns | -0.024** |

\*\*\* p < 0.001, \*\* p < 0.01, \* p < 0.05, ns = not significant

**Table S5:** Relationship of phylogenetic GoM topics with environmental variables.

| Environmental Variable | 1 | 2 | 3 | 4 |
|---|---|---|---|---|
| Soil pH | 0.031ns | -0.006ns | -0.014ns | -0.022ns |
| Bulk density | 0.014ns | -0.017* | -0.003ns | 0.003ns |
| Soil nitrogen | 0.026* | -0.008ns | -0.010ns | -0.009ns |
| Annual mean temp. (Bio1) | 0.002ns | -0.017ns | 0.019ns | -0.003ns |
| Temp. seasonality (Bio4) | -0.066ns | 0.029ns | 0.011ns | 0.019ns |
| Max. temp. of warmest month (Bio5) | 0.002ns | -0.028ns | -0.043ns | 0.067* |
| Precip. seasonality (Bio15) | -0.032ns | 0.003ns | -0.005ns | 0.028* |

\*\*\* p < 0.001, \*\* p < 0.01, \* p < 0.05, ns = not significant

**Figure S1:**
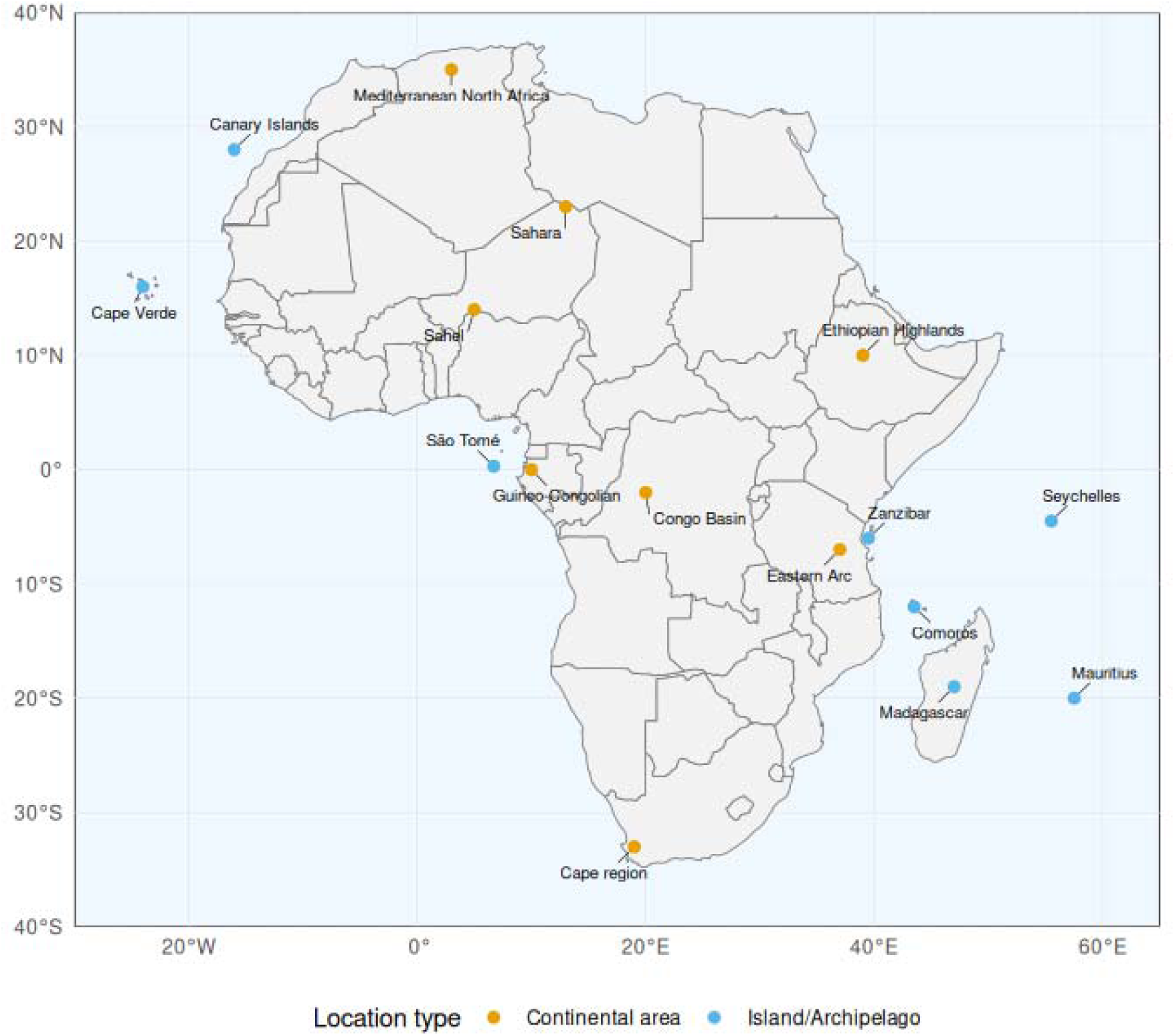
Map of Africa showing centroids of major locations mentioned in the text.

**Figure S2:**
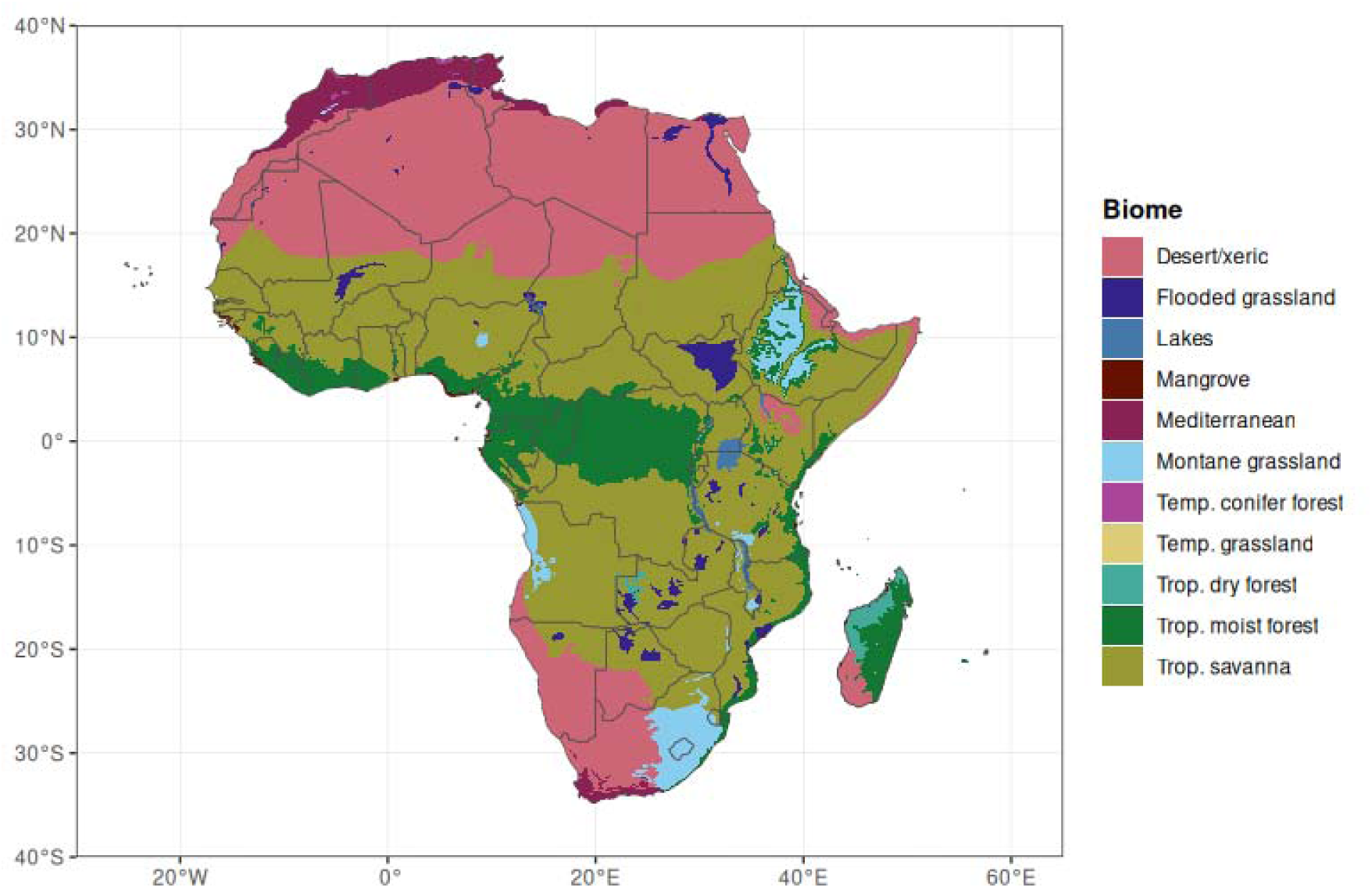
Map of African biomes. Colors correspond to biome categories used throughout this study. Biome data from Olson et al. (2001), downloaded from https://africa-knowledge-platform.ec.europa.eu/dataset/biomes.

**Figure S3:**
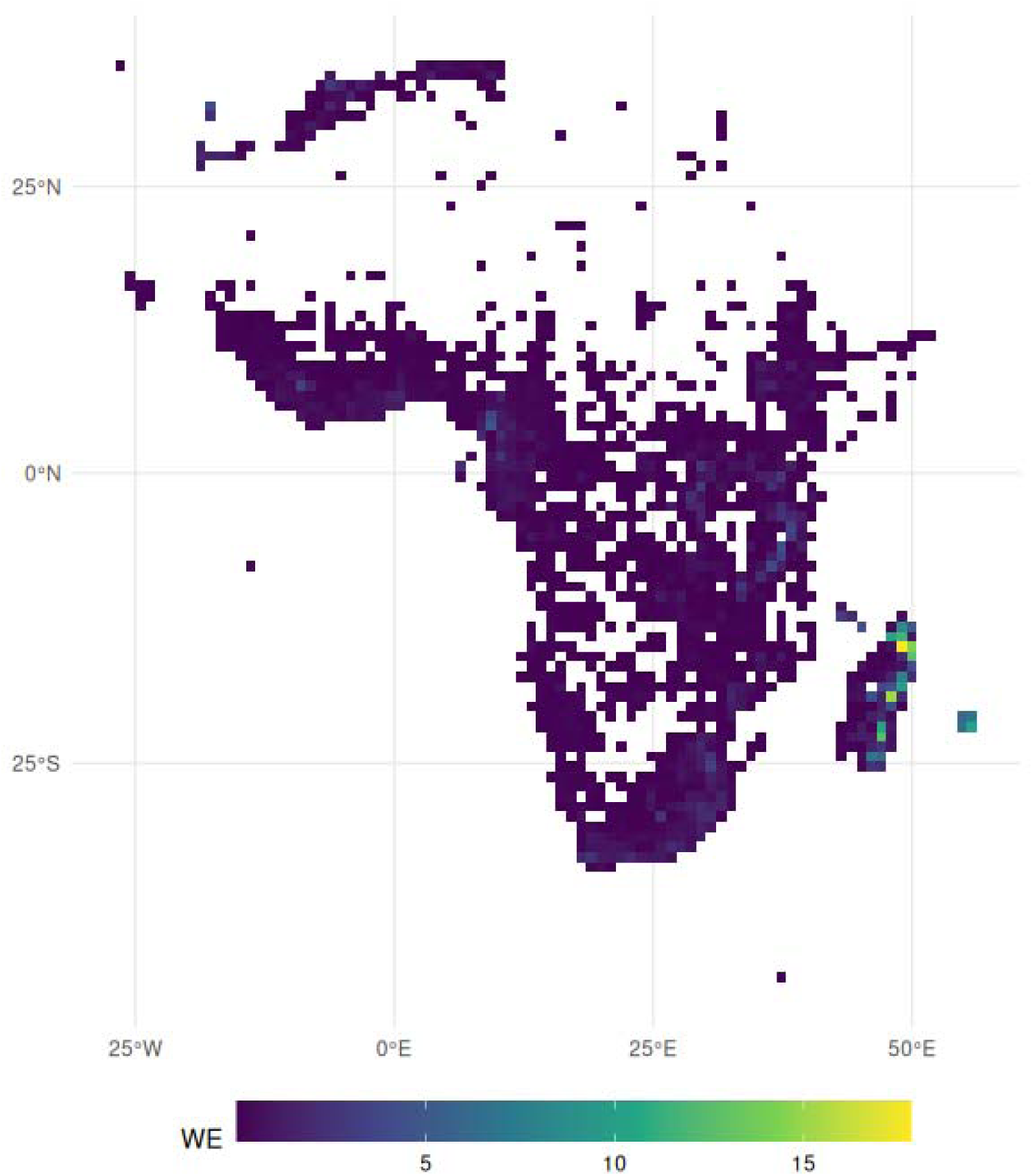
Observed weighted endemism (WE) of African ferns.

**Figure S4:**
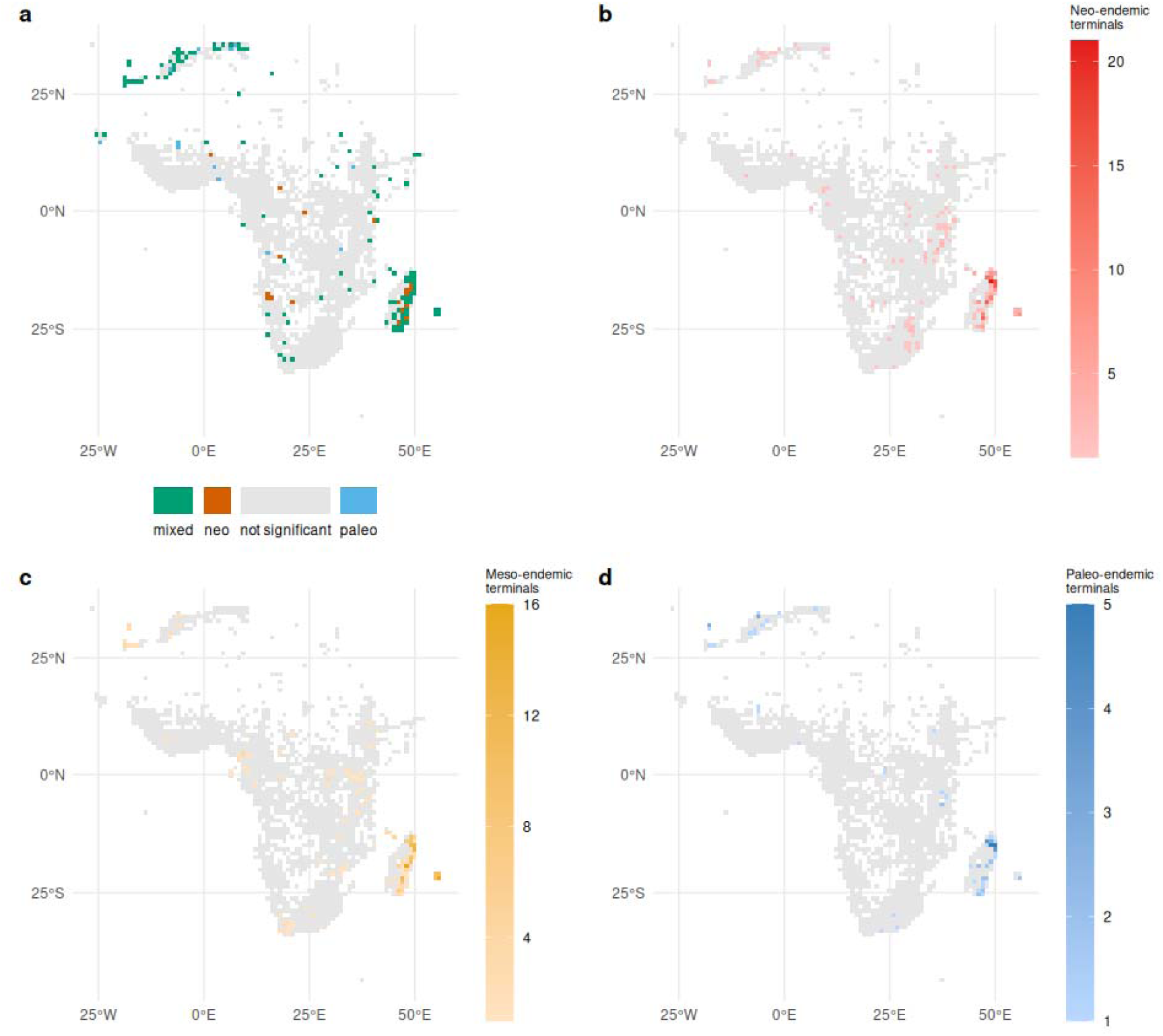
RWiBaLD analysis in relation to CANAPE results for African ferns. (a) CANAPE endemism classification. (b–d) Number of RWiBaLD-classified terminal branches per grid cell for (b) neo-endemic (red), (c) meso-endemic (yellow), and (d) paleo-endemic (blue) terminals.

**Figure S5:**
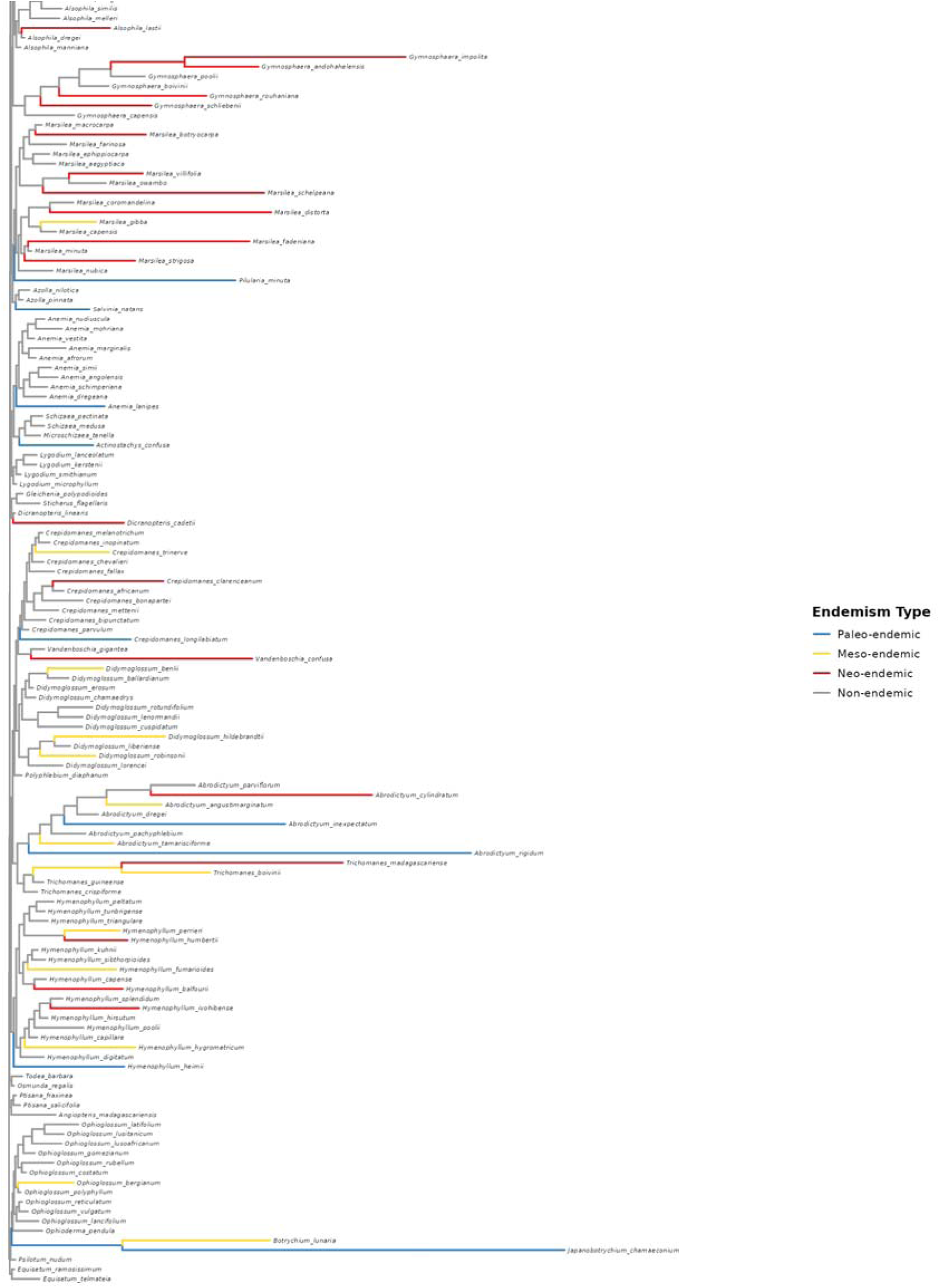
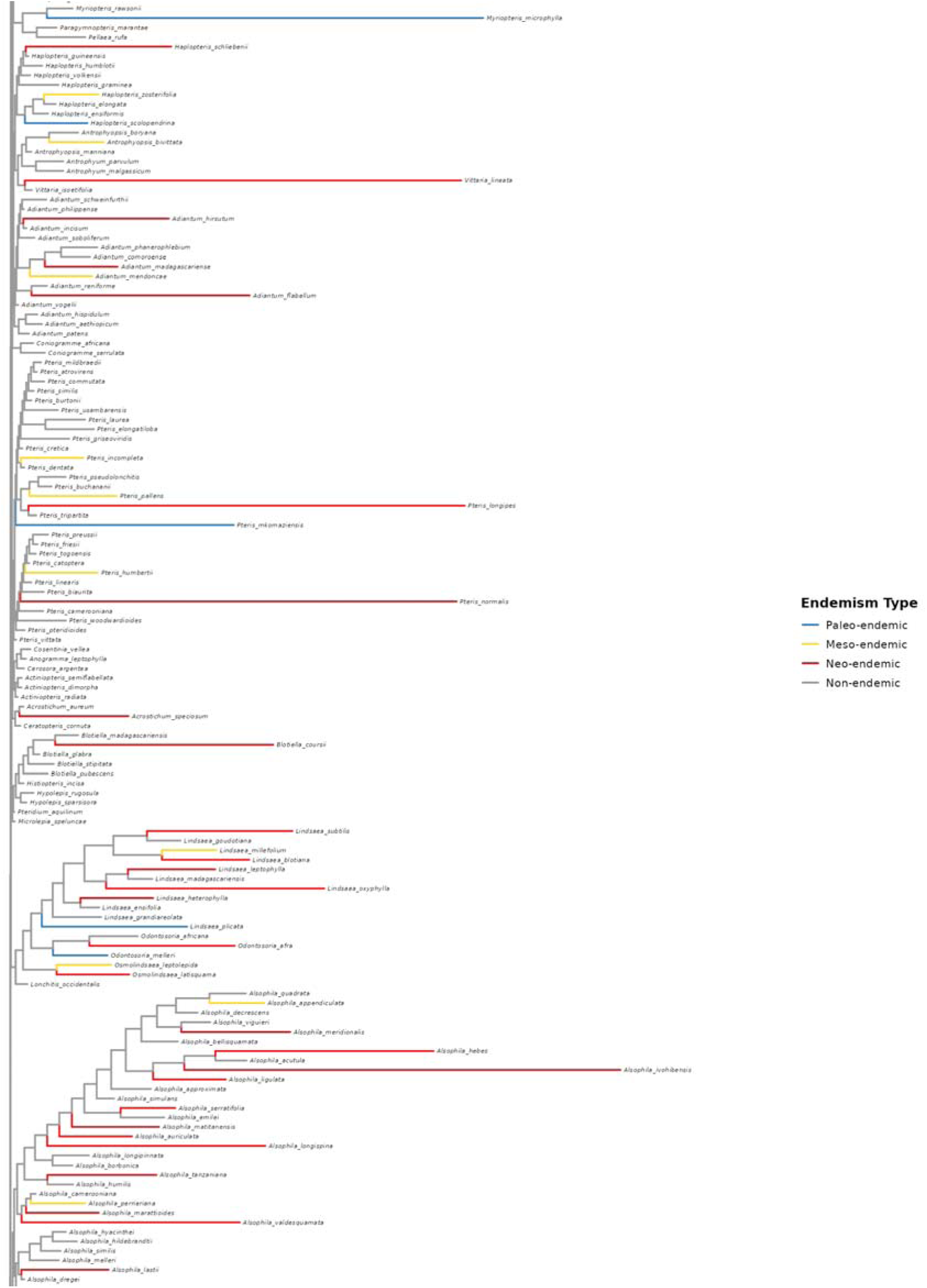
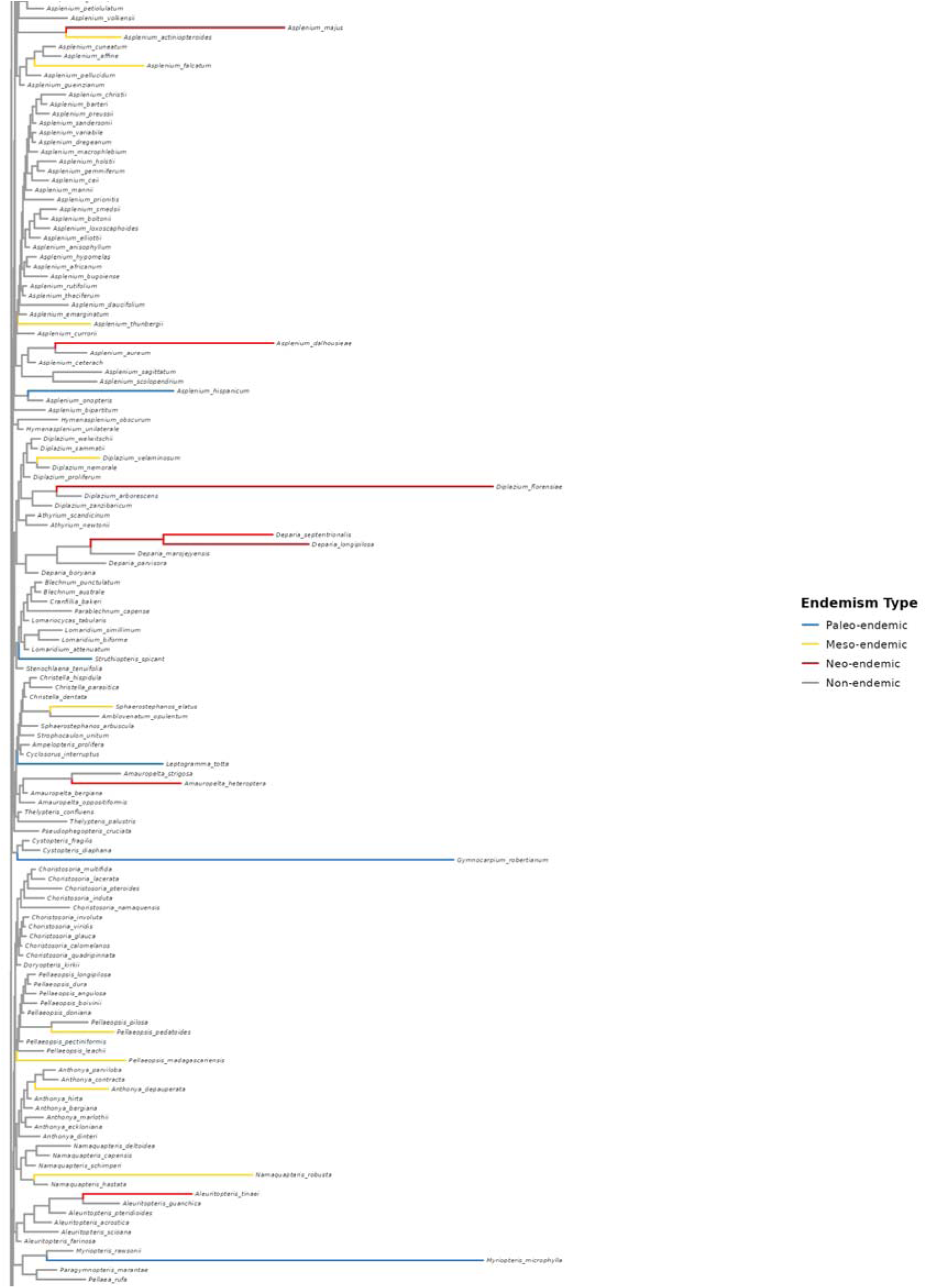
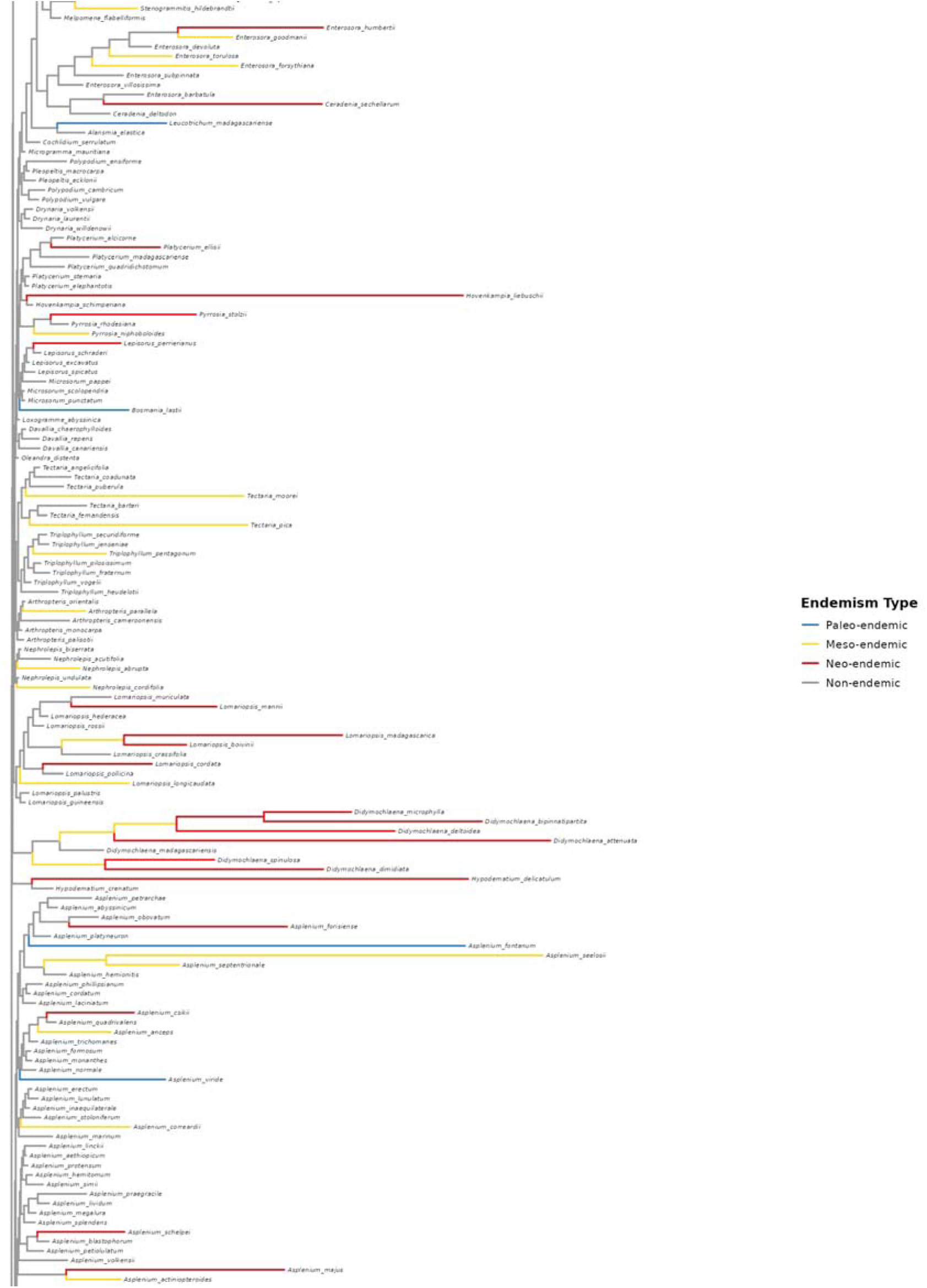
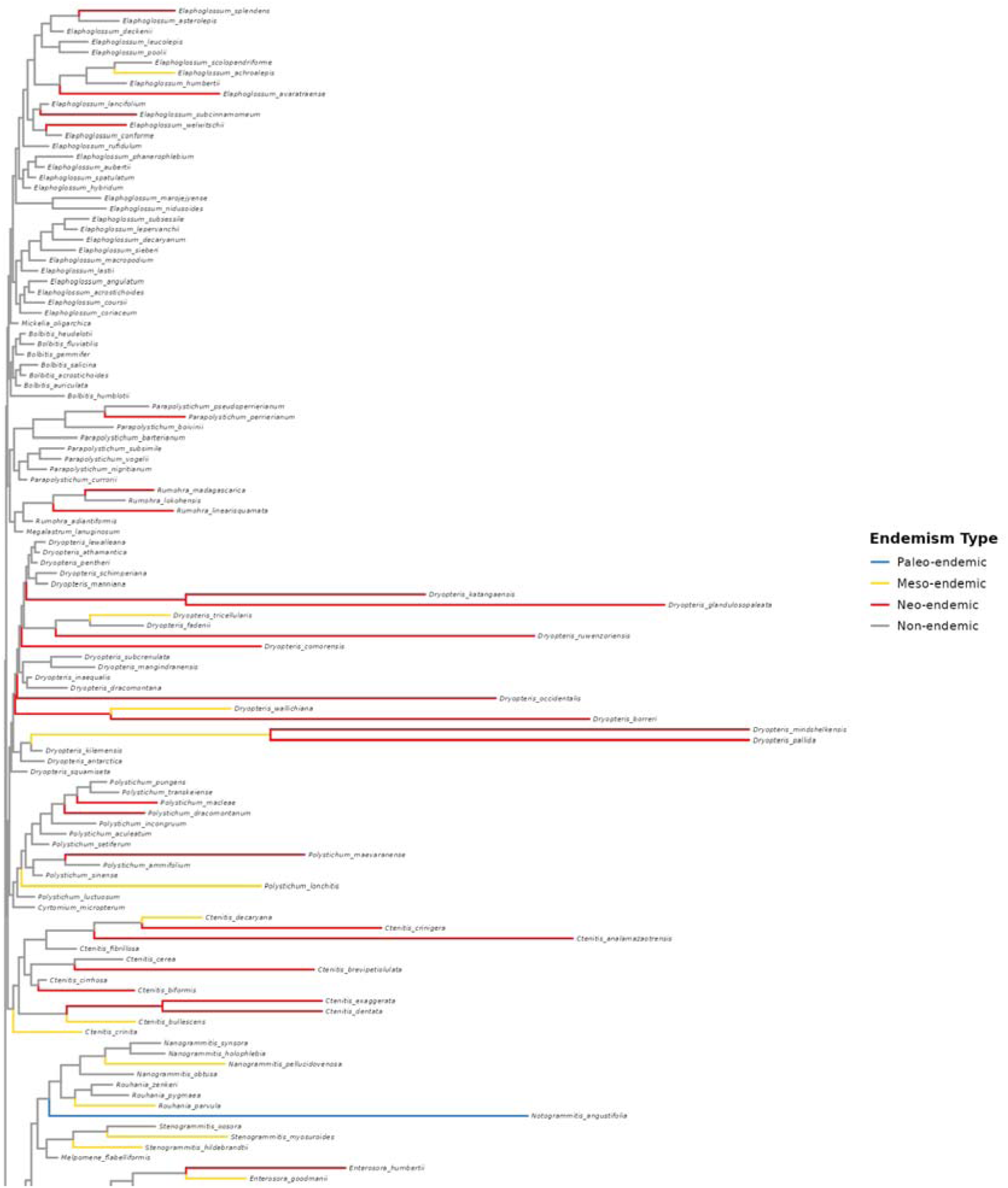
RWiBaLD scores mapped onto the range-weighted comparison tree for all branches. Branches are colored by endemism type. Figure continues on following pages.

**Figure S6:**
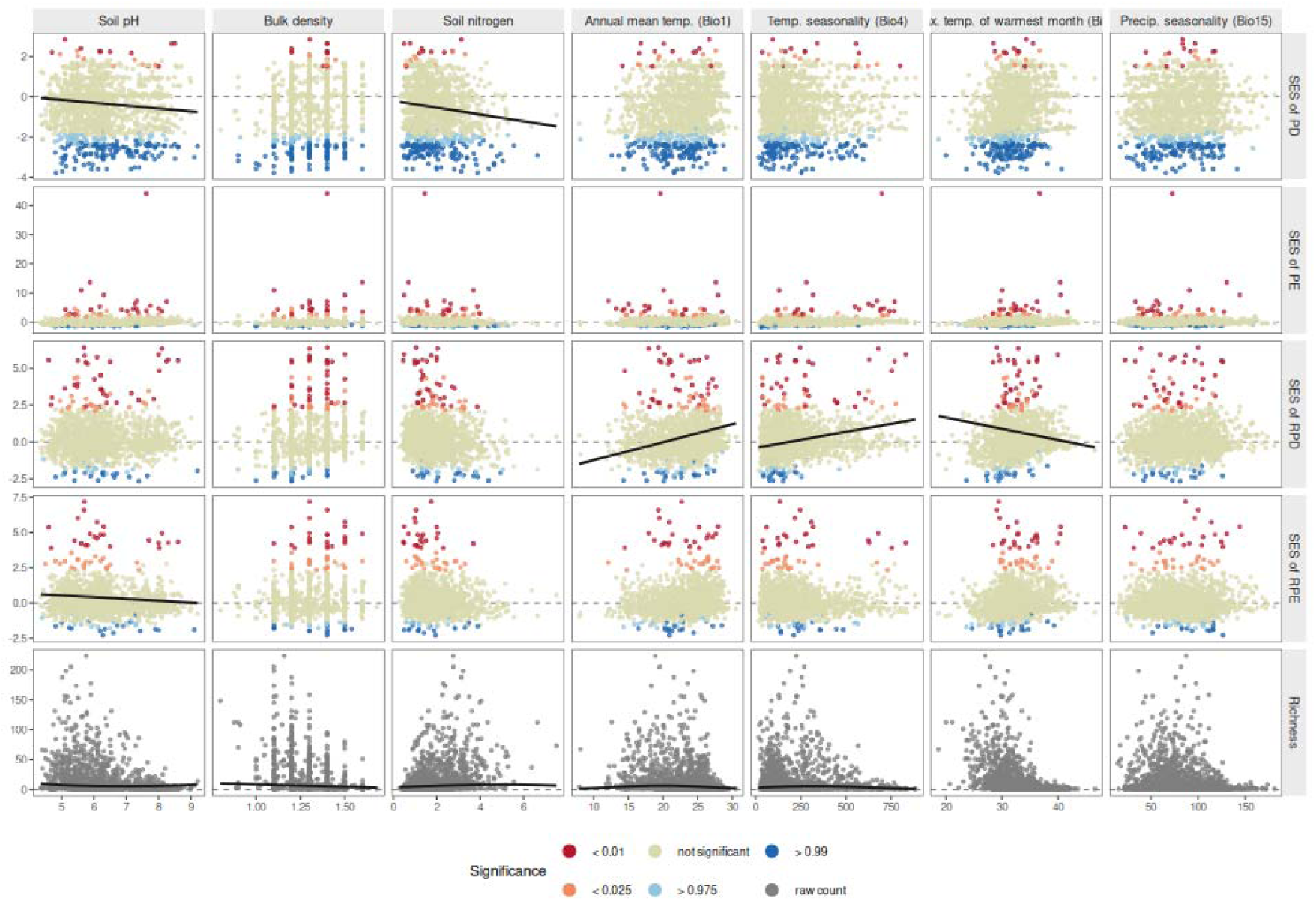
Relationship between environmental variables and alpha diversity metrics across grid cells. Each point is one grid cell. Lines indicate marginal fixed effect of significant predictors from spatial models (p < 0.05). Rows = diversity metric; columns = environmental variable. p-values (two-tailed test) of alpha diversity metrics other than species richness indicated by color.

